# Somatostatin slows Aβ plaque deposition in aged *APP^NL-F/NL-F^* mice by blocking Aβ aggregation in a neprilysin-independent manner

**DOI:** 10.1101/2022.09.26.509540

**Authors:** Declan Williams, Bei Qi Yan, Hansen Wang, Logine Negm, Christopher Sackmann, Claire Verkuyl, Vanessa Rezai-Stevens, Shehab Eid, Christine Sato, Joel C. Watts, Holger Wille, Gerold Schmitt-Ulms

**Author notes:** Please address correspondence to, Tanz Centre for Research in Neurodegenerative Diseases, University of Toronto, Rm 6KD447, 60 Leonard Ave, Toronto, Ontario M5T 0S8, Canada. Abbreviations: AD, Alzheimer’s disease; Aβ, amyloid beta; APP, amyloid precursor protein; Cort, cortistatin; IDE, insulin degrading enzyme; Mme, neprilysin gene; Ppia, peptidylprolyl isomerase A; Sst, somatostatin; Sstr, somatostatin receptor.

## Abstract

The molecular underpinnings that govern the endoproteolytic release of the amyloid beta peptide (Aβ) from the amyloid precursor protein (APP) are now quite well understood. The same cannot be said for the events that precipitate the aggregation and amyloid deposition of Aβ in Alzheimer’s disease (AD). The 14-amino-acid cyclic neuroendocrine peptide somatostatin (SST-14) has long been thought of as playing a role, foremost by controlling the expression of the Aβ clearing enzyme neprilysin, and more recently by directly interacting with Aβ oligomers. Missing have been *in vivo* data in a relevant Aβ amyloidosis model. Here we addressed this shortcoming by crossing *App^NL-F/NL-F^* mice with *Sst*-deficient mice of identical genetic background to assess if and how the presence of Sst influences key pathological hallmarks of Aβ amyloidosis that develop in *App^NL-F/NL-F^* mice after 10 months of age. Surprisingly, we found that Sst had no influence on whole brain neprilysin transcript, protein or activity levels, an observation that cannot be accounted for by a compensatory upregulation of the Sst paralog, cortistatin (Cort), that we observed in 15-month-old Sst-deficient mice. The absence of Sst did lead to a subtle but significant increase in the density of cortical Aβ amyloid plaques. Follow-on western blot analyses of whole brain extracts indicated that Sst interferes with early steps of Aβ assembly that manifest in *Sst* null brains through the appearance of SDS-stable smears of 55- 150 kDa. As expected, no effect of Sst on tau steady-state levels or its phosphorylation were observed. Results from this study are easier reconciled with an emerging body of data that point toward Sst affecting Aβ amyloid plaque formation through direct interference with Aβ aggregation rather than through its effects on neprilysin expression.

## INTRODUCTION

The peptide hormone somatostatin (SST) was initially discovered as an inhibitor of growth hormone secretion (1, 2). It is now understood that SST, along with its paralog cortistatin (CORT) (3), modulates a host of activities in the gastrointestinal, nephritic, immune, and central nervous systems (4). Both hormones are released from larger proproteins exceeding one hundred amino acids in length through endoproteolytic cleavages of C-terminal peptides which cyclize by disulfide bonding. Alternative endoproteolytic processing produces prominent variants named according to their amino acid sequence lengths as SST-14, SST-28, CORT-17, and CORT-29. These peptides have similar yet distinct physicochemical properties and affinities for five human somatostatin receptors (SSTRs), which they bind to following their release into the extracellular space (5–8). The distinct binding profiles of SST and CORT, in turn, elicit distinct physiological responses that point toward overlapping but non-identical functions (9–11).

Separate lines of biochemical, histological, and genetic evidence implicate the somatostatinergic system in Alzheimer’s disease (AD) (12–14). In fact, among the first biochemical analyses of postmortem AD brains were two studies that drew attention to a profound reduction in somatostatin levels relative to age-matched control brains (15, 16), a finding that has since been validated in human postmortem brains (17) and an AD mouse model (18).

Somatostatin-expressing neurons or interneurons may be more vulnerable to certain AD-associated stressors. For instance, cortical SST immunoreactive neurons were reported to be more susceptible to degeneration than neighboring neuropeptide Y expressing cells, with the loss of these neurons proportional to amyloid plaque and neurofibrillary tangle burden (15, 19). Similarly, somatostatinergic interneurons of the temporal cortex, but not the hippocampus or entorhinal cortex, are disproportionately lost in AD (20), compared to parvalbumin interneurons, which resist Aβ-induced toxicity in AD patients and mouse models (20, 21).

One way to rationalize these observations is to note that amyloid plaques and distressed somatostatinergic neurons are often found in close proximity, pointing toward a scenario of somatostatinergic neurons increasing the propensity for Aβ to form aggregates nearby (22, 23). A possible causal relationship has also been raised by genome-wide association studies. More specifically, the same single-nucleotide polymorphism in the 3’ untranslated region of the *SST* gene was associated with increased late-onset AD risk in two genetically separate population cohorts in Finland and China, particularly among individuals carrying the apolipoprotein E-ε4 allele (24, 25).

What might be the mechanism through which SST modulates AD? SST was initially reported to promote Aβ clearance indirectly. Thus, in mouse cortical neuronal cultures, a modest decrease in extracellular Aβ_42_, but not Aβ_40_, followed a single administration of nanomolar to low micromolar SST-14. This effect was attributed to an SST receptor-dependent transcriptional activation of neprilysin (26), a protease that the same group had previously shown to digest Aβ (27). SST has also been shown to enhance the proteolytic activity of insulin degrading enzyme (IDE) toward Aβ *in vitro*, to increase microglial IDE transcription and translation, as well as phagocytosis of Aβ (28), and to be an IDE substrate itself (29, 30).

More recently, a direct influence of SST on Aβ emerged when we identified SST by mass spectrometry as the smallest human protein that interacts selectively with oligomeric but not monomeric Aβ (31). We then established that nanomolar concentrations of SST or CORT modulate the aggregation of Aβ_42_ (31). Since then, the selective SST binding to oligomeric Aβ_42_ has been shown to extend to a β-sheet pore-forming Aβ_42_ tetramer with putative neurotoxicity (32). The possibility that SST may modulate Aβ plaque formation directly is heightened by the fact that this molecule spontaneously forms amyloid fibrils *in vitro* (33) and is, in fact, stored prior to its synaptic release in dense granules as a natural amyloid (34). Upon their release, it takes in the order of minutes to hours before these SST amyloids are dissolved (35), providing an opportunity to influence nearby Aβ aggregation in AD (12).

Remarkably, despite all these observations implicating somatostatin as a possible factor in AD spanning decades, no prior experimental study has investigated the influence of SST in an *in vivo* Aβ amyloidosis paradigm. Here, we set out to begin to address this unmet need by studying intercrosses between *Sst* null mice and a humanized *App* knock-in mouse line that is known to develop a profound Aβ amyloidosis as the mice age. Breeding the cross as *Sst* hemizygotes provided progeny of mixed *Sst* genotype which consistently developed Aβ amyloid plaques from middle to late adulthood. We present data that queried how *Sst* deficiency affected (i) transcript levels of cortistatin and neprilysin, (ii) Aβ amyloid plaque burden and plaque distribution, (iii) neprilysin protein and activity levels, (iv) SDS-stable Aβ oligomers, as well as (v) steady-state tau levels and tau phosphorylation. To generate these data, we made use of *App^NL-F/NL-F^Sst^-/-^* mice, with age-matched Sst-expressing *App^NL-F/NL-F^Sst^+/+^* mice serving as controls.

## MATERIALS AND METHODS

### Mouse models

*App^NL-F^* mice (36) were generously provided by Drs. Takashi Saito and Takaomi C. Saido of the Laboratory for Proteolytic Neuroscience, RIKEN Brain Science Institute, Hirosawa, Wako-shi, Saitama, Japan. The B6N.129S4(129S6)-Sst^tm1Ute^/J Sst knockout mouse line (37) was supplied by the Jackson Laboratory (catalog number 008117, Bar Harbor, ME, USA).

Double transgenic animals of all three *Sst* genotypes were housed together with no more than 5 animals per cage. Water was offered to the mice *ad libitum* and their feed was 18% protein chow. The cages were kept at elevated room temperature (24°C) in an environment that was subjected to an artificial twelve-hour day and night cycle. Daily health checks for activity and overall appearance of the mice were conducted. The cages were changed once a week. No behavioral studies were undertaken. At 12 and 15 months of age, the mice were sacrificed exclusively for brain collection. To this end, the mice were deeply anesthetized by isoflurane inhalation and exsanguinated by two-minute transcardiac perfusion with phosphate buffered saline. Upon extraction, each brain was separated at the midline; then the right and left hemispheres transferred to neutral buffered 10% formalin (Sigma-Aldrich, St. Louis, MO, USA) and dry ice, respectively. Right cerebral hemispheres were transferred to 70% ethanol (v/v) after 48 hours of fixation, left hemispheres were stored at -80°C. Littermates of different *Sst* genotypes were sampled wherever possible.

### Genotyping

Genomic DNA was isolated from tail clips collected at weaning. After overnight Proteinase K (Bioshop, Burlington, ON, Canada) digestion at 55°C, genomic DNA was isolated by phenol extraction then precipitated and purified with ethanol. The following primers were used to inform the mouse breeding program and to determine *Sst* genotypes of experimental animals: *App* wild-type forward (5’-ATCTCGGAAGTGAAGATG-3’) and reverse (5’-GTTAAGTTCTCATCTACA- 3’), *Sst* wild-type (5’-TCAGTTTCTGCAGAAGTCTCTGGC-3’) and knockout (5’- ATCCAGGAAACCAGCAGCGGCTAT-3’) forward and Sst reverse (5’- GAATGCCAATAGTTTGCGCAGCTCC-3’). *Sst* genotyping was performed in accordance with the supplier’s protocol. A second forward primer used to confirm the *Sst* deficient allele was 5’- AGGCACGATGGCCGCTTTGG-3’.

### Antibodies

Monoclonal 4G8 anti-APP/Aβ, applied at 1:2000 dilution (catalog number 800701) and monoclonal 6E10 anti-APP/Aβ, applied at 1:1000 (catalog number 803001) were from BioLegend (San Diego, CA, USA). Monoclonal 82E1 anti-Aβ, applied at 1:1000 dilution was from Immuno-Biological Laboratories (catalog number 10323, Minneapolis, MN, USA). Monoclonal Tau5 anti-Tau (catalog number AHB0042), applied at 1:5000 dilution, monoclonal AT8 anti- Ser202/Thr205-phospho-Tau (catalog number MN1020), applied at 1:1000 dilution, and AT180 anti-Thr231-phospho-Tau (catalog number MN1040), applied at 1:1000 dilution, were sourced from Thermo Fisher Scientific (Mississauga, ON). Anti-neprilysin (catalog number AF1126), applied at 1:1000 dilution, was from Bio-Techne (Minneapolis, MN, USA). Horseradish peroxidase-conjugated horse anti-mouse IgG (catalog number 7076), applied at 1:5000 dilution, was from Cell Signaling Technology (Danvers, MA, USA).

### Western blot analyses

Frozen left cerebral hemispheres of mice were weighed and homogenized with 0.7 mm diameter zirconia beads (catalog number 11079107, BioSpec Products, Bartlesville, OK, USA) in a buffer composed of 100 mM Tris-HCl, pH 8.3, 100 mM NaCl, 2x Roche PhosSTOP phosphatase inhibitor cocktail (catalog number 4906837001, Roche, Basel, Switzerland), 2x Roche cOmplete protease inhibitor cocktail (catalog number 11836170001, Roche) with three 1-minute pulses of bead-beating and 1 minute of cooling on ice between each pulse. Note that detergent was omitted during homogenization to minimize foaming but was added for the solubilization of membrane proteins during the subsequent protein extraction. Specifically, homogenates were extracted with 1% NP40, 100 mM Tris-HCl, pH 8.3, 100 mM NaCl, 1x Roche PhosSTOP phosphatase inhibitor cocktail, 1x Roche cOmplete protease inhibitor cocktail, at 4 °C for 30 minutes with agitation before centrifugation at 4 °C for 5 minutes at 2000 g and 15 minutes at 21130 g. Supernatants of brain protein extracts were collected following centrifugation, then protein concentrations were determined by bicinchoninic acid assay (catalog number 23227, Thermo Fisher Scientific) and equalized by dilution in extraction buffer. After samples were prepared with NuPAGE LDS Sample Buffer (catalog number NP0007, Thermo Fisher Scientific) containing 2% β-mercaptoethanol, and heated at 70°C for 10 minutes, 30 to 60 µg of protein per lane were loaded and separated on 10% Bis-tris BOLT SDS-Page gels (catalog number NW00105BOX, Thermo Fisher Scientific) at 100V for 1.5 hours in MOPS buffer. Proteins were then transferred to 0.45 µm pore PVDF (catalog number IPVH00010, Millipore) at 25 to 35 V for 1 hour and incubated with appropriate antibodies overnight at 4 °C, followed by 1-hour incubation with Horseradish-Peroxidase-linked secondary antibodies at room temperature.

Horseradish peroxidase-catalyzed chemiluminescence from SuperSignal West Dura Extended Duration HRP Substrate (catalog number 37071, Thermo Fisher Scientific) was detected on radiography film and band intensities were quantified on ImageJ software version 1.53e.

### Neprilysin activity measurements

To assess neprilysin activity, brain samples were analyzed using the Sensolyte 520 Neprilysin Activity Assay Fluorometric Kit (catalog number AS-72223, AnaSpec) and following the manufacturer’s instructions. Briefly, 10% brain extracts were generated and cleared of insoluble material as described in the western blot analysis section. Throughout these steps, the brain extracts were kept on ice. Next, 50 µL of brain extract supernatants were pipetted onto a 96- well black/clear bottom plate (catalog number 165305, Thermo Fisher Scientific) and 50 µL of 2x Neprilysin Substrate (provided in the kit) was added to the samples for an assay concentration of 5 µM substrate. The 96-well plate was then placed in a 37°C incubator for 1 hour followed by fluorometric analysis using a SpectraMax i3x Multi-Mode Microplate Reader (Molecular Devices, San Jose, CA, USA) at 490nm/520nm excitation/emission wavelengths. As positive controls and to calibrate the system served recombinant neprilysin enzyme (provided in the kit and diluted to assay concentrations of 1 to 10 ng per well). Background activities toward the Neprilysin Substrate were determined by undertaking assay reactions in the presence of 100 nM levels of the neprilysin inhibitor thiorphan (provided in the kit). All analyses were undertaken in triplicate.

### Immunohistochemistry

Brain hemispheres were paraffin-embedded then sectioned parasagittally at 5-micron thickness, and three contiguous sections per animal from near lateral 0.1 mm were mounted to each slide. Tissue processing was based on the following steps: xylene deparaffinization, hydration in graded ethanol/water, peroxidase inactivation in methanolic 3% H_2_O_2_ (v/v) for 25 minutes, antigen retrieval in formic acid for 5 minutes, blocking with horse serum (catalog number MP-7802-15, Vector Laboratories, Burlingame, CA, USA), overnight 4G8 (1:3000) antibody application in mouse-on-mouse diluent (catalog number BMK-2202, Vector Laboratories). Staining of 4G8 immunoreactivity was with 3,3’-diaminobenzidine. Nuclei were stained with Harris hematoxylin (Sigma-Aldrich) then hydrochloric acid was used to differentiate the tissue.

Prepared slides were scanned at 20x magnification (0.5 micrometers/pixel) on an Aperio Scanscope AT2 (Leica, Wetzlar, Germany) and plaque quantification was performed on HALO software (Version 2.3.2089.29, Indica labs, Albuquerque, NM, USA). Margins of the cortical and hippocampal regions were manually defined (as exemplified in **Fig 3B)**, creating two annotation layers per slide. The anterior olfactory nucleus and corpus callosum were included in the cortical annotation. Vasculature and meninges were manually excluded from the annotations. Classifiers for plaques (4G8-positive tissue), non-plaque (4G8-negative tissue) and background (voids in the tissue) were defined manually and iteratively to compensate for differences in background staining such that the minimum number of classifiers were applied to the dataset. Following automated detection of the classifiers, the quantified annotations of each slide were manually examined to verify that the classifier settings discriminated plaque, non-plaque, and background. Raw data (object data) from each analysis were exported in text format. The areas of all objects classified as plaques and non-plaques from a given annotation on a given slide were summed to give the total cortical or hippocampal area, respectively.

### Reverse transcription real-time quantitative polymerase chain reaction (RT-qPCR)

Working surfaces were treated with RNase Zap (catalog number AM9780, Thermo Fisher Scientific). Frozen left cerebral hemispheres were weighed then homogenized in QIAzol lysis reagent (catalog number 79306, Qiagen, Venlo, Netherlands) on ice with an immersion style tissue homogenizer for 40 seconds, then RNA was extracted and purified with an RNeasy kit (catalog number 74004, Qiagen) according to the manufacturer’s protocol. RNA integrity and yield were assessed by 1.5% agarose gel electrophoresis and UV-visible spectroscopy (NanoDrop, Thermo Fisher Scientific), respectively. An AffinityScript cDNA synthesis kit (catalog number 200436, Agilent, Santa Clara, CA, USA) was used according to the manufacturer’s instructions to prepare cDNA libraries.

TaqMan™ Universal Master Mix II and FAM^TM^-minor groove binder-labeled TaqMan probes (Cort: Mm00432631_m1, Mme: Mm00485028_m1, Ppia: Mm02342430_g1, Sst: Mm00436671_m1, Thermo Fisher Scientific) were mixed with cDNA in the manufacturer’s recommended volumetric ratios. A total of 180 ng of cDNA was used in each reaction to quantify all transcripts except *Mme*, for which 50 ng was used. Fluorescence curves were obtained on a Light Cycler 480 II thermal cycler (Roche) in runs starting with a 2-minute 50°C hold, then a 10-minute 95°C hold, then 40 cycles from 95°C for 15 seconds to 60°C for 1 minute when fluorescence was measured at 465-510 nm. For each *App^NL-F/NL-F^*genotype and each transcript studied, analyses were based on a minimum of three biological replicates. Quantification cycles (C_q_), also known as threshold cycles (C_t_), were determined using the second derivative maximum method. Sample-matched C_q_ values generated by the concomitant analysis of *Ppia* on each plate were used as references to calculate relative transcript abundances based on the delta-delta C_t_ method (38).

### Neprilysin Activity Assay

To assess neprilysin activity in the brain extracts of 15-month-old *App^NL-F/NL-F^* used for immunoblotting analysis, samples were analyzed using the Sensolyte 520 Neprilysin Activity Assay Fluorometric Kit (catalog number AS-72223, AnaSpec, Fremont, CA, USA) that employs a 5-FAM/QXL FRET substrate. Upon incubation with a sample of interest comprising neprilysin activity the 5-FAM component is freed from the substrate and can be quantified fluorometrically. Briefly, the 10% (w/v) brain extracts were centrifuged at 15,000 rcf for 5 minutes to sediment insoluble debris. 50 uL aliquots of the centrifuge supernatants were then pipetted into the wells of a 96-well black, flat-bottom plate (catalog number 165305, Thermo Fisher Scientific) along with 50 uL of 2x 5-FAM/QXL FRET substrate. A total of four brain extracts each from wildtype and *Sst* knockout *App^NL-F/NL-F^* mice were analyzed, with each sample being tested in three technical replicates. As a positive control served the FAM/QXL FRET substrate incubated with 10 ug/mL recombinant neprilysin provided in the kit. For the generation of negative background fluorescence controls, the substrate was replaced with water in the assay mix. The 96-well plate was then placed in a 37°C incubator for 1 hour followed by fluorometric analysis using a SpectraMax i3x Multi-Mode Microplate Reader at 490 nm excitation and 520 nm emission wavelengths and the activity was determined in relative fluorescence units, normalized to the recombinant neprilysin samples.

### Statistics

All processing of raw data and all statistical tests were undertaken with Microsoft Excel and GraphPad. This included RT-qPCR analyses collected with LightCycler 480 (version 1.5.1.62), immunohistochemistry data obtained with HALO software (Indica Labs, Albuquerque, New Mexico, USA), western blots scanned for densitometry analyses with ImageJ (National Institutes of Health and the Laboratory for Optical and Computational Instrumentation, University of Wisconsin, WI, USA), and neprilysin activity analyses. Specifically, the Microsoft Excel Analysis ToolPack and the GraphPad *t*-test calculator were used to compute *p-*values. All statistical analyses made use of the two-tailed *t*-test based on the assumption that the variance of groups of samples was not known. Amyloid plaque image analysis data from cohorts of age- and Sst genotype-matched mice were assessed for normality using kurtosis and skewness values determined with the descriptive statistics function, where acceptable values for either metric were between -2 and 2. Plaque density distributions were considered discrete. *p*-values are either shown as numbers or with asterisks according to conventions in the field, i.e., *p*-values lower than 0.05 were considered significant and were marked with a single asterisk. Additional asterisks are shown for every tenfold lower *p*-value. When *t*-tests failed to meet the significance threshold, the abbreviation ‘ns’ is shown.

### Ethics statement

All mouse handling procedures were in accordance with the Canadian Council on Animal Care, reviewed and approved by the University Health Network Animal Care Committee under Animal Use Protocol 4183. The use of mouse brains for biochemical applications detailed in this report was reviewed and authorized (Biosafety Permit 208-S06-2) by the Environmental Health a Safety office at the University of Toronto, Toronto, Ontario, Canada.

## RESULTS

### Generation of *App^NL-F/NL-F^Sst^-/-^*mouse line

Both the *Sst* and the *App* genes are coded on Chromosome 16 in mice, which could jeopardize the ability to generate mice carrying both the *App* gene mutation and *Sst* deletion through intercrossing if the genes were linked and recombination disfavored. A closer analysis mostly dispelled this concern because the *Sst* gene maps to cytoband qB1 (23889573-23890958 bp, Genome Reference Consortium mouse build 38/mm10) and the *App* gene is coded in cytoband qC3.3 (84952666-85175255 bp), located 32 cM apart (**Fig 1A**).

**Figure 1.**
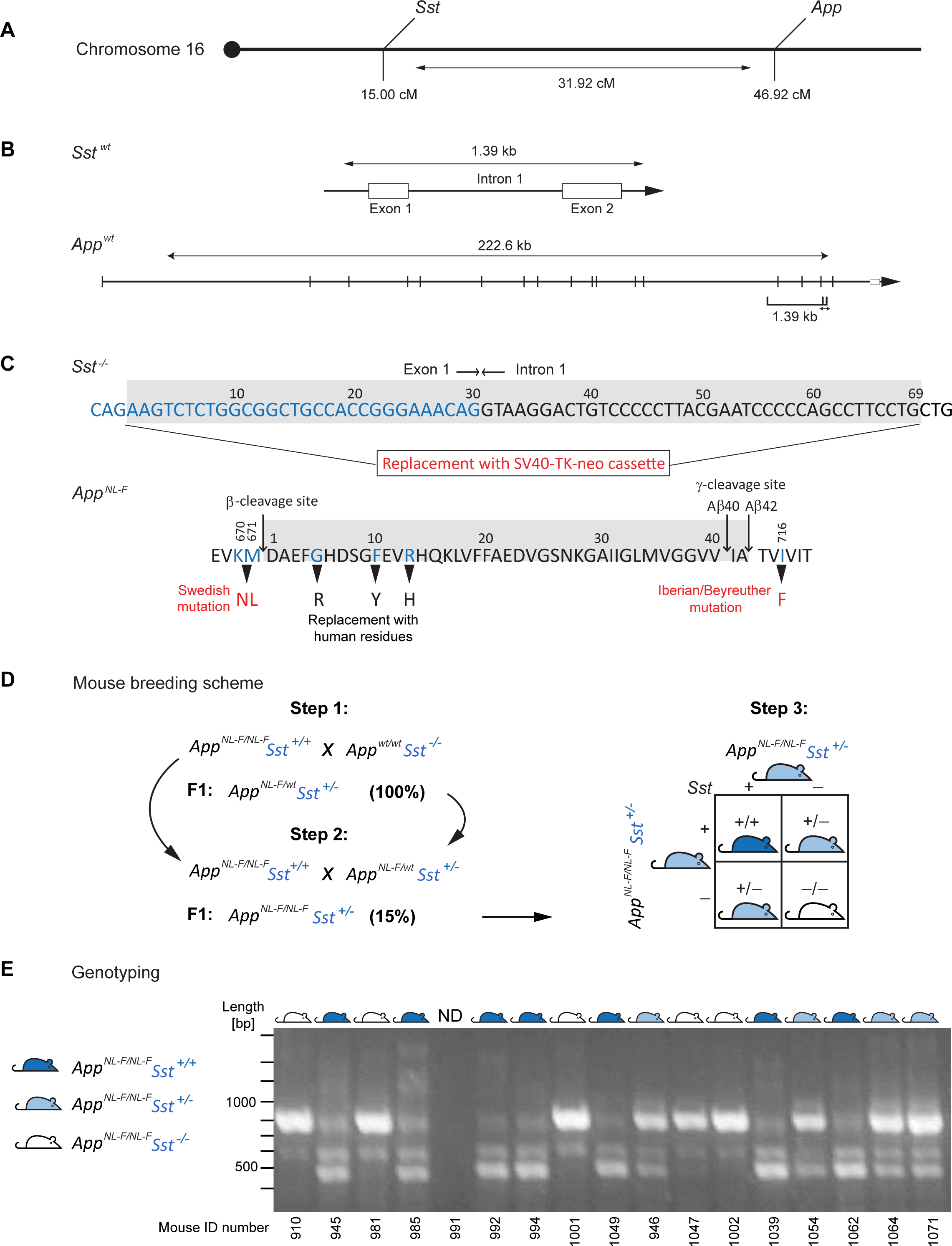
Generation of the *App^NL-F/NL-F^Sst^-/-^* mouse line. (A) Depiction of the relative positioning of *Sst* and *App* genes on mouse Chromosome 16. (B) Intron-exon gene organizations of mouse *Sst* and *App* genes. (C) Summary of genomic alterations present in the *Sst^-/-^* and *App^NL-F/NL-F^* parental mouse lines used in this study. (D) Depiction of the three-step breeding scheme, including Punnett Square diagram of final intercrosses. (E) Representative results from tail clip-based genotyping analyses of *App^NL-F/NL-F^*X *Sst^+/-^* intercrosses.

Sst and App proteins are coded by multi-exon gene sequences (**Fig 1B**). Several mice in which the *Sst* gene or the expression of Sst receptors were deleted have been reported in the literature (39). Here we employed a well-validated *Sst* null line that was originally generated by the insertion of a SV40-TK-neo cassette, which had caused the deletion of 69 base pairs within the Exon 1-Intron 1 junctional sequence region within the *Sst* gene (Zeyda *et al.*, 2001) (**Fig 1C**). To our knowledge the sequence of this insert was not originally published, which is why we inserted it here as a supplement (**S1 Fig**). This *Sst* null line was selected because it had been reported to feature normal hippocampal App expression. Moreover, the aforementioned data, which suggested Sst-deficiency leads to diminished neprilysin activity and elevated Aβ_42_ levels, were also generated with this line (26).

The selection of a compatible Aβ amyloidosis model fell on a previously reported knock- in mouse line sharing with the *Sst* null mice the same C57BL/6 inbred genomic background (36).

This *App^NL-F/NL-F^* line carries several mutations that flank or alter the Aβ42 coding sequence. Cumulatively these changes humanize Aβ, increase production of Aβ42 three-fold, and drive the Aβ42-to-Aβ40 brain production ratio more than 10 times higher, relative to wild-type (36). We chose this knock-in model to avoid insertion artefacts and variances that are frequently observed in transgenic lines due to the large and fluctuating number of transgene copies. Prior global proteome analyses of whole brain and hippocampus samples from these mice indicated that Sst expression is comparable in *APP^NL-F/NL-F^* and APP^wt/wt^ control mice (Cort was not detected in these analyses) (40, 41).

To generate cohorts for this study, three consecutive intercrosses were undertaken. More specifically, we initially crossbred the *Sst* null line with homozygous *App^NL-F/NL-F^* mice (**Fig 1D**). Next, the F1 generation (*App^NL-F/wt^Sst^+/-^*) was backcrossed with *APP^NL-F/NL-F^* mice to produce intercrosses that were homozygous for the *NL-F/NL-F* mutation but heterozygous for the Sst knockout (*App^NL-F/NL-F^Sst^+/-^*). Due to the need for a Chromosome 16 crossover event to achieve this genotype only approximately 15% of the F1 offspring from this second intercrossing, i.e., 13 out of a total of 87 mice, were confirmed as *App^NL-F/NL-F^Sst^+/-^*. Finally, *App^NL-F/NL-F^Sst^+/-^*were intercrossed and progeny were genotyped from tail clippings using a customized PCR analysis (**Fig 1E**). This scheme minimized undesired variability by ensuring that *Sst* null mice (*App^NL-F/NL-F^Sst^-/-^*) shared the same parents and housing as their wild-type *Sst* littermates (*App^NL-F/NL-F^Sst^+/+^*). Genotyping validated that *App^NL-F/NL-F^Sst^-/-^*, *App^NL-F/NL-F^Sst^+/-^*, and *App^NL-F/NL-F^Sst^+/+^*progeny were obtained in approximate Mendelian ratios. *App^NL-F/NL-F^Sst^+/-^*hemizygotes and *App^NL-F/NL-F^Sst^-/-^* knockouts were physically and behaviorally unremarkable and lived past 16 months of age with a mortality equivalent to their *Sst* wild-type littermates.

### *Sst*, *Mme*, and *Cort* transcript expression in *App^NL-F/NL-F^Sst^-/-^*mouse brains

Prior work indicated that mRNA transcript levels of the *Sst* paralog *Cort* may undergo a gender-dependent compensatory increase in *Sst* knockout mice (42). Moreover, transcription of the neprilysin gene (*Mme*) has been reported to be induced by Sst (26), a finding that could have direct implications for interpreting effects of Sst on Aβ amyloid deposition, due to the enzyme-substrate relationship between neprilysin and Aβ. To explore these possibilities, we aged *Sst* wild-type, hemizygous, or knockout *App^NL-F/NL-F^* mice to 12 or 15 months, ages at which the *App^NL-F/NL-F^* genotype gives rise to subtle and prominent Aβ amyloid deposition respectively (**Fig 2A**). We then determined transcript levels of *Sst*, *Mme*, *Cort*, and *Ppia*, the latter a common proxy for total RNA in RT-qPCR of brain tissue. As biological source materials served mid-sagittal cut half brains, as opposed to specific brain regions, to minimize variances that can be introduced during dissection steps.

**FIGURE 2.**
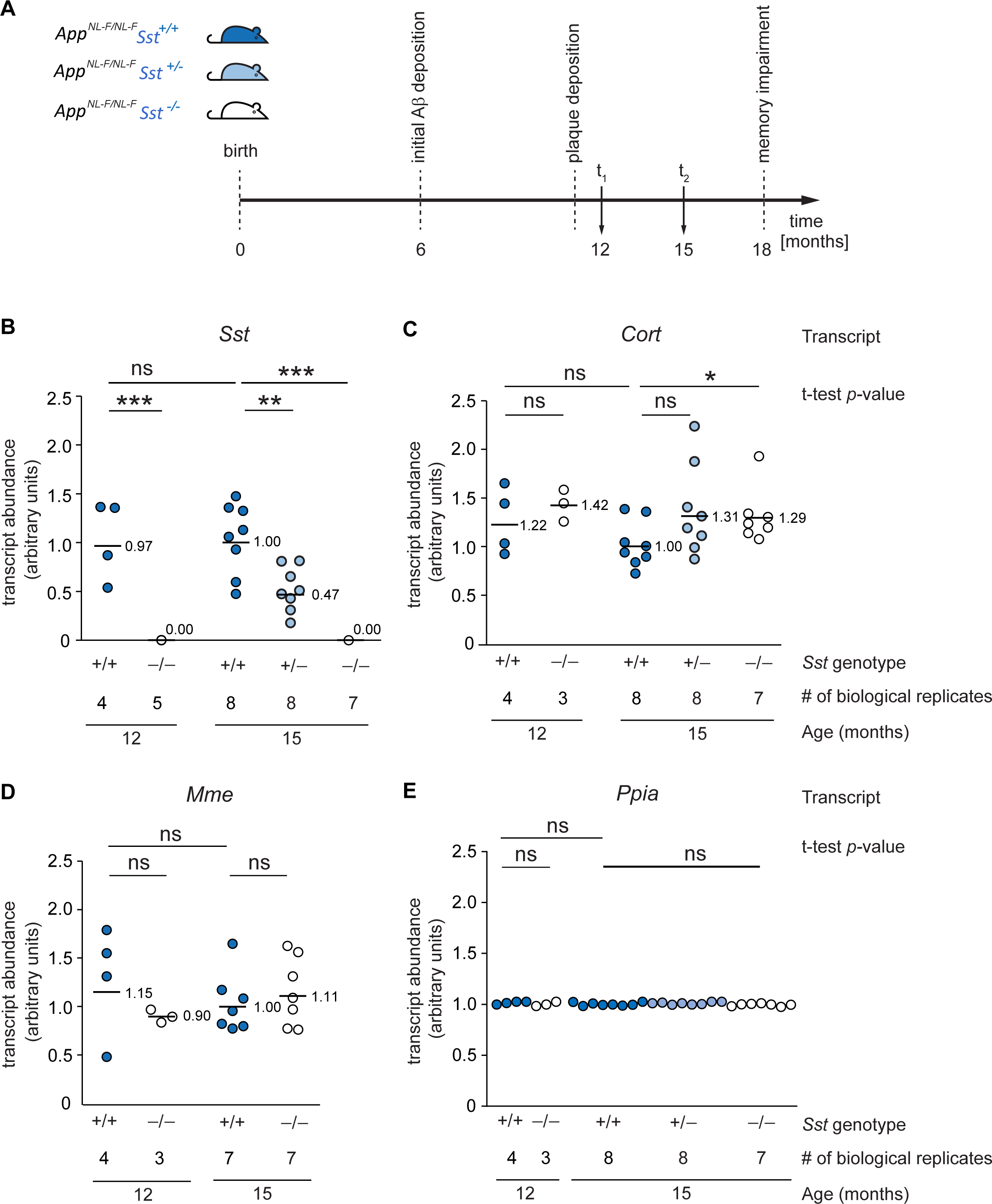
*<u>Sst</u>*-deficient *App^NL-F/NL-F^Sst^-/-^* mice exhibit increased cortistatin and unaltered neprilysin transcript levels relative to *App^NL-F/NL-F^Sst^+/+^* mice. (A) Timeline of relevant pathological events and ages selected for comparative RT-qPCR and neuropathological analyses. At times t_1_ and t_2_, mouse brains were bisected in the mid-sagittal plane. One half provided source material for RT-qPCR and western blot analyses, the other half was fixed in formalin and assessed for Aβ deposition by immunohistochemistry. (B) *Sst* transcript levels in whole brain extracts of *App^NL-F/NL-F^Sst^+/+^*, *App^NL-F/NL-F^Sst^+/-^*, and *App^NL-F/NL-F^Sst^-/-^* mice. *Sst* transcript levels were lower in 15-month-old hemizygotes than in wild-types (p = 0.004, paired two-tailed t-test) and were undetectable in 12- or 15-month-old *Sst* knockouts. (C) Brain *Cort* transcript levels did not differ significantly between *Sst* wild-type and hemizygous mice (paired two tailed t-test) but were significantly increased in 15-month-old *Sst* knockout *App^NL-F/NL-F^Sst^-/-^* mice, relative to *Sst* wild-type *App^NL-F/NL-F^Sst^+/+^*mice. (D) Whole brain neprilysin (*Mme*) transcript levels were not affected by mono- or bi-allelic *Sst* knockout in *App^NL-F/NL-F^Sst^-/-^* mice. (E) Relative *Ppia* mRNA levels in the brains of 12- and 15-month-old *App^NL-F/NL-F^Sst^+/+^*, *App^NL-F/NL-F^Sst^+/-^*, and *App^NL-F/NL-F^Sst^-/-^* mice. Each dot is the value from one mouse. *Ppia* transcription was assumed to be constant in all *Sst* genotypes. All amplifications shown in subpanels of this figure were based on 180 ng cDNA. Bars are averages.

Distributions of transcript abundance for *Sst* (**Fig 2B**), *Cort (***Fig 2C**), and *Mme* (**Fig 2D**) were approximately normal, and coefficients of variation ranged from 0.35 to 0.43, 0.12 to 0.34, and 0.07 to 0.44, respectively. Moreover, for all transcripts measured, mRNA levels in males and females were equivalent, as assessed by *t*-test. As expected, these analyses revealed that *Ppia* transcript levels were highly consistent between all samples (**Fig 2E**), indicating that both the neurophysiological backgrounds sampled, and the sample preparations were consistent.

The brain *Sst* mRNA concentration of *App^NL-F/NL-F^Sst^+/-^*hemizygotes averaged around half that of *Sst* wild-type *App^NL-F/NL-F^Sst^+/+^*animals, suggesting that in most hemizygote animals a compensatory transcriptional upregulation of the remaining *Sst* allele did not occur (**Fig 2B**).

However, overlapping distributions of *Sst* transcript levels in *Sst* wild-type *App^NL-F/NL-F^Sst^+/+^* and *Sst* hemizygote *App^NL-F/NL-F^Sst^+/-^*mice indicated a degree of flexibility of *Sst* expression and raised the possibility that some hemizygous individuals may produce sufficient levels of *Sst* to be phenotypically wild-type. The *Sst* transcript was undetectable in all *App^NL-F/NL-F^Sst^-/-^* mice, confirming that the mutated allele was not expressed.

At both ages examined, *Sst* wild-type *App^NL-F/NL-F^Sst^+/+^* mice had consistent levels of *Cort*, *Mme*, and *Sst* transcripts, indicating that each of these genes was stably expressed over the 3 month age range (**Fig 2B-D**). *Cort* mRNA transcript levels increased in *Sst*-deficient *App^NL-F/NL-F^Sst^-/-^*mice by approximately one third (35%), relative to age-matched *Sst* wild-type *App^NL-F/NL-F^Sst^+/+^* mice. This increase was significant (*p*<0.05) between *Sst* wild-type and knockout *App^NL-F/NL-F^* cohorts at 15 months of age (**Fig 2C**). Remarkably, there was no significant difference in neprilysin transcript levels between *Sst* wild-type *App^NL-F/NL-F^Sst^+/+^* and *Sst* knockout *App^NL-F/NL-F^Sst^-/-^* animals at 12 or 15 months of age (**Fig 2D**).

In summary, these RT-qPCR data validated a compensatory increase in *Cort* mRNA levels in response to *Sst* knockout but did not corroborate the previously reported male sex-specificity of this increase (42). These data also did not validate the concept of Sst acting as an inducer of whole brain neprilysin expression.

### Amyloid plaque deposition in *App^NL-F/NL-F^Sst^-/-^*mice

The RT-qPCR data may lead to an anticipation that the compensatory increase in *Cort* transcript levels could diminish effects of *Sst* knockout on Aβ deposition. To address this question experimentally, we next assessed cohorts of *App^NL-F/NL-F^*Sst^+/+^, *App^NL-F/NL-F^Sst^+/-^*and *App^NL-F/NL-F^Sst^-/-^* mice for Aβ deposition at 12 and 15 months of age. To this end, half brains were formalin fixed, paraffin-embedded and parasagittally cut. Next, 4G8 anti-Aβ immunoreactivity was visualized with the horseradish peroxidase substrate 3, 3-diaminobenzidine, and Aβ plaque densities were quantified (**Fig 3A**). Immunoreactivity was confined mainly to the cortex, hippocampus, and olfactory bulb (**Fig 3B-D**). A small number of amyloid plaques were occasionally observed within the corpus callosum and the anterior olfactory nucleus, particularly in brain sections of 15-month-old mice. The brainstem, midbrain, and cerebellum were devoid of 4G8 immunoreactivity in all samples. Of all cortical areas, the prelimbic and orbital areas generally had the lowest plaque density. Cortical plaque density increased from 12 to 15 months of age in each *Sst* genotype (*App^NL-F/NL-F^Sst^+/+^*: 3.7 times; *App^NL-F/NL-F^Sst^+/-^*: 4.6 times; *App^NL-F/NL-F^Sst^-/-^*: 4.5 times). At both ages studied, the sex of the animals did not appear to influence the propensity or speed of plaque formation that we captured in these analyses.

**Figure 3.**
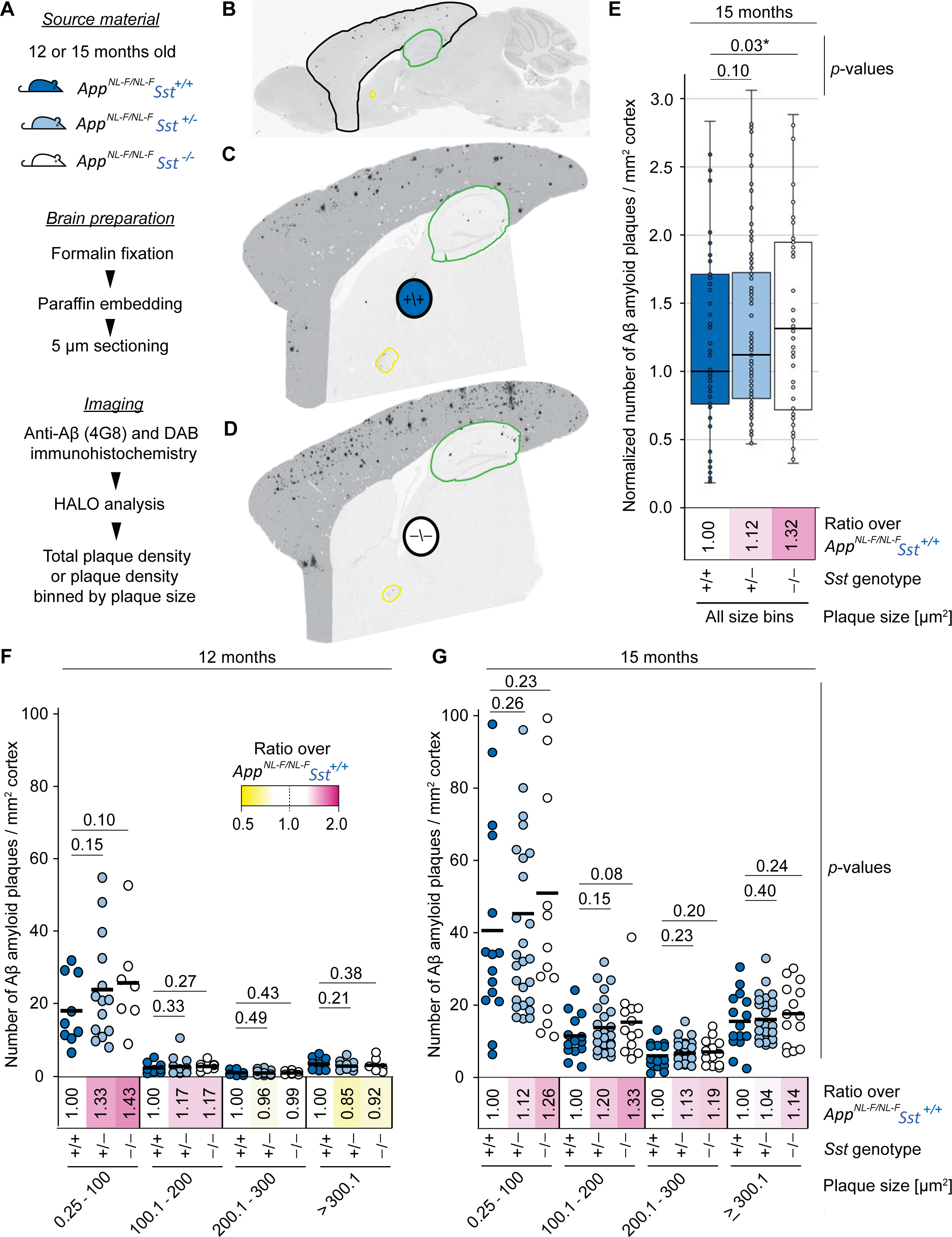
Cortical Aβ plaque density is increased in *App^NL-F/NL-F^* mice upon *Sst* ablation. (A) Workflow for the assessment of Aβ plaque distribution and density. (B) Parasagittal brain section highlighting cortical and hippocampal areas that were separately analyzed for plaque density and are shown bounded in black and green, respectively. The anterior commissure is outlined in yellow. Representative micrographs of parasagittal brain sections from 15-month- old (C) *App^NL-F/NL-F^Sst^+/+^*and (D) *App^NL-F/NL-F^Sst^-/-^* mice stained with 4G8-DAB showing plaques are largely restricted to cortical and hippocampal areas. (E) Plaque density measurements were based on all plaques exceeding 0.25 µm and were normalized relative to the median plaque density observed in *App^NL-F/NL-F^Sst^+/+^*cortices. Paired two-tailed t-test results indicate a significant relative increase in plaque density in *App^NL-F/NL-F^Sst^-/-^* cortices relative to side-by-side processed cortices derived from age-matched *App^NL-F/NL-F^Sst^+/+^* mice. (F) Comparison of cortical Aβ amyloid plaque densities, binned into four Aβ amyloid plaque size ranges, in 12-month and (G) 15-month old *App^NL-F/NL-F^* mice with wild-type, hemizygous, or knockout *Sst* genotype. Aβ plaque densities increased in all four plaque size categories in the older animals, irrespective of *Sst* genotype. A consistent trend, indicating an increase in plaque density, yet not meeting significance conventions (*p*<0.05), was observed in all four plaque size ranges in 15-month-old mice, when *Sst* hemizygous or deficient mice were compared against wild-type *Sst* expressing *_App_NL-F/NL-F* _mice._

When comparing the cortical Aβ amyloid plaque density for all plaques larger than 0.25 µM in diameter (the smallest plaque size that could be reliably detected in these analyses), we observed that *Sst*-deficient *App^NL-F/NL-F^Sst^-/-^*mice exhibited a statistically significant increase in plaque density, relative to *Sst* wild-type *App^NL-F/NL-F^Sst^+/+^* at 15 months (**Fig 3E**).

To dig deeper into how *Sst* deficiency might affect Aβ amyloid deposition, we segregated Aβ plaques with diameters up to 300 µM into three equally proportioned size ranges. Perhaps not surprisingly, the smallest plaque size range contained the largest proportion of total plaques in all three mouse populations at both ages. We were particularly interested in the relative densities of amyloid plaques of the smallest size in 12-month-old mice, hypothesizing that *Sst* influences the earliest steps of Aβ aggregation. The *Sst*-associated difference in plaque densities was indeed most pronounced at this relatively early timepoint in the size range that included the smallest detectable plaques (0.25-100 µm) but did not reach statistical significance (**Fig 3F**). In the second smallest size range examined (100-200 µm), *Sst*- deficient mice had 17% more plaques than Sst wild-types. Larger plaques (>200 µm) made up a smaller proportion of the total plaque area, namely 16%, 12%, and 11% of total plaques in *Sst* wild-type, hemizygous and deficient *App^NL-F/NL-F^* mice, respectively, and appeared equally abundant in 12-month-old mice in all three *Sst* genotypes. At 15 months, *Sst* knockout *App^NL-F/NL-F^Sst^-/-^*had consistently higher cortical plaque burdens than *Sst* wild-type mice across all plaque sizes, and hemizygote *App^NL-F/NL-F^Sst^+/-^*animals presented with intermediate plaque burdens (**Fig 3G**). Statistical analyses confirmed these trends yet did not establish significance for any of the pair-wise comparisons.

Hippocampal amyloid plaque density in 12- and 15-month-old *App^NL-F/NL-F^*mice of all three *Sst* genotypes was considerably lower than that of the cortical regions. As in the cortex, hippocampal plaques up to 200 µm were more abundant in 12-month-old Sst-deficient mice and this trend extended to all size categories at 15 months of age (**S2 Fig**).

Taken together, these analyses corroborated the notion that the presence of a functional *Sst* gene slows the formation of Aβ amyloid plaque deposition in *APP^NL-F/NL-F^* mice.

### Steady-state protein levels and activity of neprilysin in mouse brain extracts

Although we did not observe Sst-dependent differences in neprilysin transcript levels, we wondered if Sst increases neprilysin steady-state protein levels. To investigate this possibility, we quantified neprilysin protein levels in NP-40 solubilized brain extracts (derived from mid-sagittal cut half brains) from 15-month-old mice by western blot analyses. Analogous to the RT-qPCR results, this experiment revealed consistent steady-state neprilysin levels and no apparent changes in post-translational modifications between *Sst* wild-type *App^NL-F/NL-F^Sst^+/+^* and knockout *App^NL-F/NL-F^Sst^-/-^* mice (**Fig 4A**). Next, we measured the *in vitro* activity of *Sst* wild-type and deficient *App^NL-F/NL-F^* brain extracts toward a commercial fluorescent neprilysin substrate (5- FAM/QXL-520). In these analyses, background (non-neprilysin) activities within brain extracts toward the fluorescent substrate were elucidated by incubating brain extracts in the presence of the neprilysin-specific inhibitor thiorpan (**Fig 4B**). To estimate total neprilysin activity levels in our extract fractions we compared their activity to the activity of a known amount of recombinant human neprilysin. These analyses established that the presence or absence of Sst did not affect neprilysin expression or activity in a significant manner in total brain extracts.

**Figure 4:**
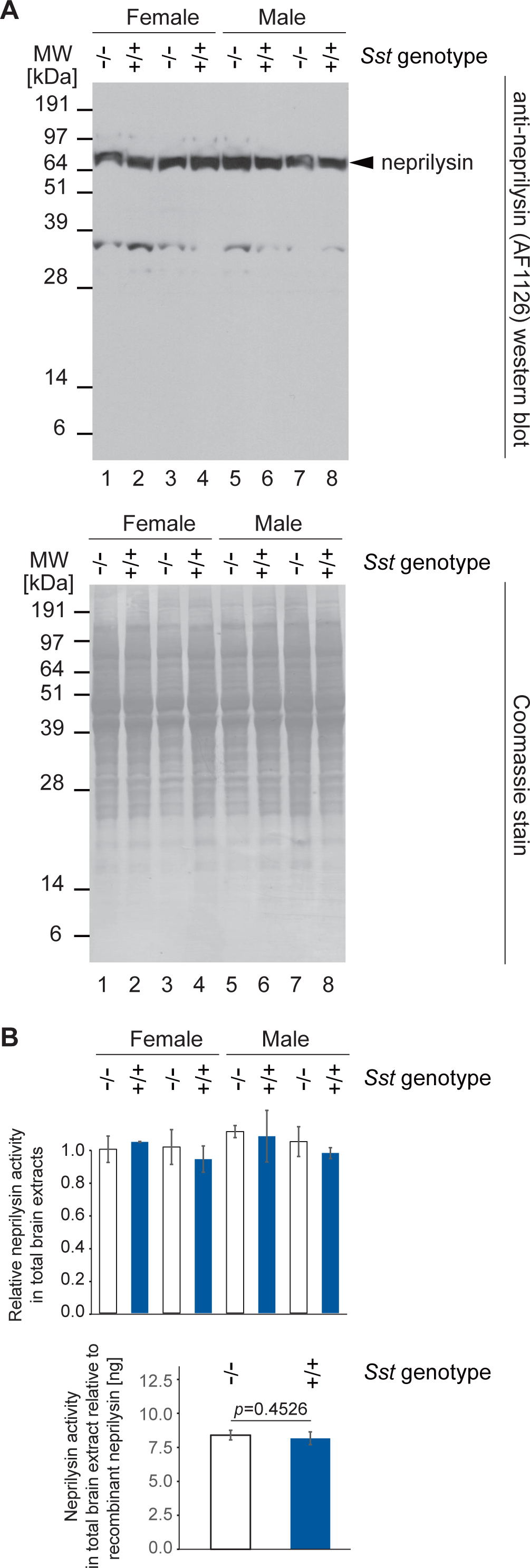
No detectable change in steady-state neprilysin protein levels or activity in Sst- deficient *App^NL-F/NL-F^*mice. (A) *Sst* gene knockout had no effect on steady-state neprilysin protein levels in extracts from mid-sagittal cut half brains of *App^NL-F/NL-F^*mice. The lower panel depicts a Coomassie stain of the SDS-PAGE-separated brain extract fractions used in these analyses whose total protein levels had been adjusted by bicinchoninic acid analysis. (B) *Sst* gene knockout had no significant effect on neprilysin activity present in brain extracts from mid-sagittal cut half brains of *App^NL-F/NL-F^* mice toward an internally quenched 5-carboxyfluorescein (5-FAM) conjugated neprilysin substrate. Whereas the upper chart shows the neprilysin activities within Sst wild-type and deficient *App^NL-F/NL-F^*brain extracts relative to each other (normalized to the mean neprilysin activity measured in one of the Sst-deficient *App^NL-F/NL-F^* mice), the lower chart depicts their mean neprilysin activities relative to the activity of a known amount of recombinant human neprilysin.

### *Sst* ablation promotes the formation of high molecular mass Aβ oligomers

We considered that Sst may influence the earliest Aβ aggregation steps, namely oligomerization. This idea was based on our earlier observation that the addition of Sst to a synthetic Aβ_1-42_ preparation delays Aβ *in vitro* aggregation in a concentration-dependent manner (31) and on data from molecular dynamics simulations by our collaborators showing that the presence of Sst interferes with early Aβ oligomer assembly (43). We made use of the monoclonal 82E1 antibody, which exclusively detects Aβ assemblies with exposed Aβ N-termini and, consequently, will not bind APP or its derivatives encompassing the noncleaved Aβ sequence (44). The same 15-month-old brain extracts analyzed by 82E1 western blot had striking differences between *Sst* wild-type *App^NL-F/NL-F^Sst^+/+^*and knockout *App^NL-F/NL-F^Sst^-/-^* mice: Whereas *Sst* knockout *App^NL-F/NL-F^Sst^-/-^* mice gave a strong 82E1-reactive smear covering apparent molecular weights of 55-150 kDa, wild-type *App^NL-F/NL-F^Sst^+/+^* gave rise to considerably weaker signals in this molecular weight range (**Fig 5A**). Densitometry showed that the difference in signal intensities in this molecular weight range were highly significant (**Fig 5B**).

**Figure 5:**
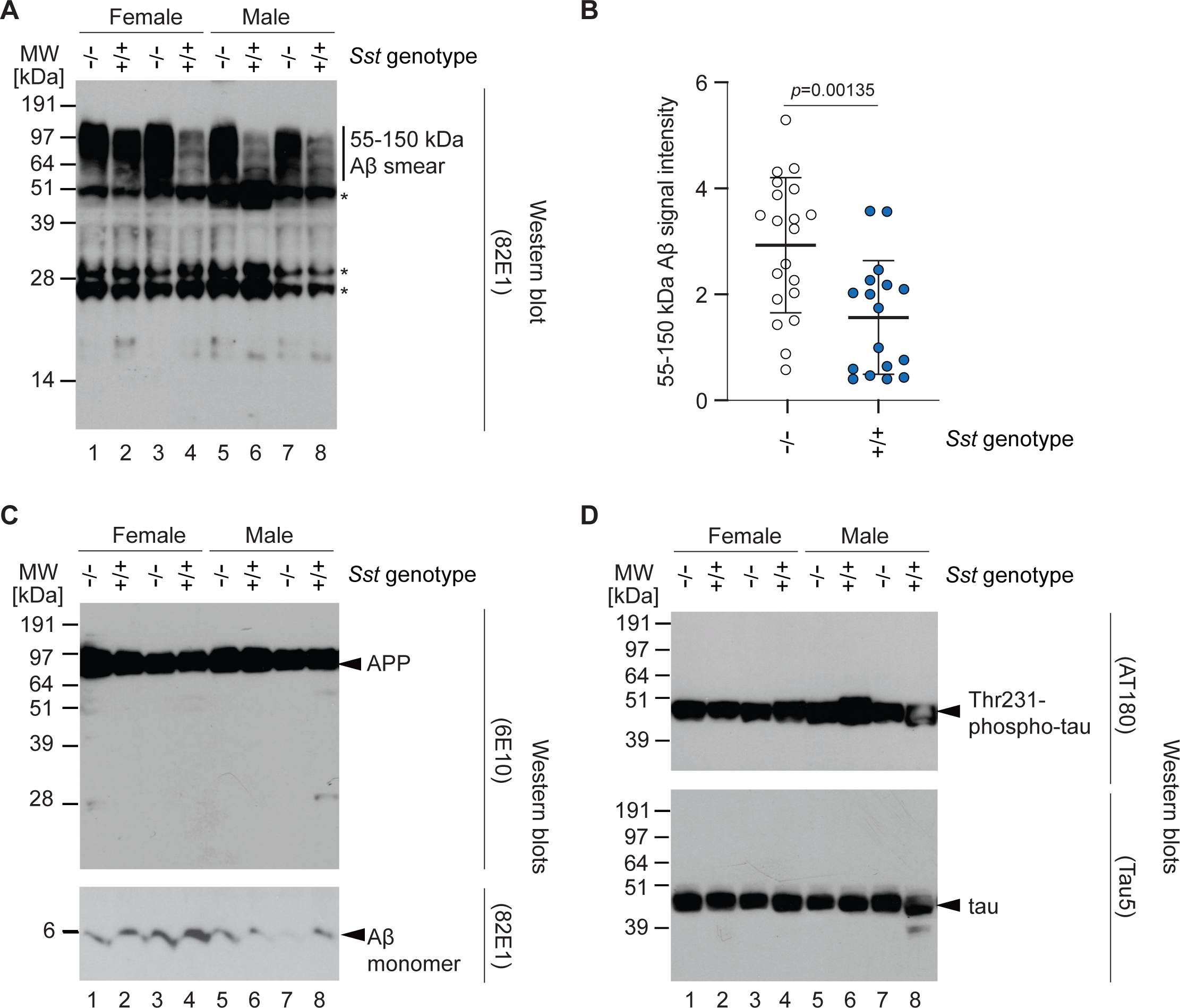
*Sst* ablation promotes the formation of high molecular mass Aβ oligomers. (A) *Sst* gene knockout is associated with an increase in the intensity of diffuse 82E1 signals observed at 55-150 kDa in western blot analyses of *App^NL-F/NL-F^* mouse-derived brain extracts. Bands indicated in this panel with asterisks represent signals generated by the secondary anti- mouse antibody reacting with endogenous IgG present in the mouse brain extracts. IgG was identified based on bands of the same size and relative intensity also appearing when the same anti-mouse secondary antibody was applied without primary antibody to brain extract fractions. (B) ImageJ-based densitometry analyses of western blot bands shown in Panel A within the 55-150 kDa apparent molecular weight range validate that the increase in signal intensities is significant (*p*=0.0052) when comparing *Sst* wild-type and deficient *App^NL-F/NL-F^* mice. (C) Germline *Sst* ablation does not translate into differences in total APP or monomeric Aβ levels in 15-month-old *App^NL-F/NL-F^*mice, detected with 6E10 and 82E1 (following long exposure) antibodies, respectively. (D) Although AT180 phospho-tau levels were subject to small fluctuations in mouse brain extracts of 15-month-old *App^NL-F/NL-F^*, these fluctuations did not track with the expression or loss of *Sst* gene expression. Similarly, no *Sst* gene associated differences in total tau levels were observed in these 15-month-old *App^NL-F/NL-F^* mice. Fractions analyzed in this figure were adjusted for total protein levels using the bicinchoninic acid method and were identical to those depicted following Coomassie staining in Figure 4A.

Next, we generated western blots with the 6E10 anti-Aβ epitope, which predominantly detects full-length APP in extracts of 15-month-old *App^NL-F/NL-F^*upon low exposure of blots, to determine if *Sst* influences the expression of this precursor. We also produced a long exposure of the low molecular weight region of the 82E1 western blot for the detection of monomeric Aβ (**Fig 5C**). Finally, we probed western blots with antibodies directed against total tau and a prominent tau phosphorylation epitope encountered in AD brain samples (**Fig 5D**). No changes in APP, monomeric Aβ, total tau or phospho-tau were observed.

Taken together, experiments in this section revealed that Sst slows Aβ deposition primarily through a neprilysin-independent effect on Aβ oligomerization that appears to affect early steps in Aβ oligomer assembly. Accordingly, Sst ablation increased the formation of SDS-resistant Aβ oligomers but did not influence levels of the APP precursor or the production of monomeric Aβ.

## DISCUSSION

This study was designed to reveal whether *Sst* gene ablation affects Aβ amyloidosis *in vivo*. To this end, we crossed the extensively studied C57BL/6-derived *Sst* null mouse line (37) and *App^NL-F/NL-F^*mice (36). The *Sst* gene knockout was validated by genotyping and RT-qPCR and was observed to cause a slight compensatory upregulation of the mRNA levels of the *Sst* paralog *Cort*. Remarkably, neither whole brain neprilysin transcript nor protein or activity levels were impacted by *Sst* genotype, yet *Sst* ablation still led to significantly stronger cortical Aβ deposition in 15-month-old *App^NL-F/NL-F^* mice. When Aβ plaques were binned by size in 12-month-old *App^NL-F/NL-F^* mice, which are at early stages of plaque deposition, it became apparent that the *Sst* gene ablation promoted the formation of the smallest Aβ plaques. A follow-on investigation into the mechanism revealed that in the absence of Sst the signal intensities of SDS-stable Aβ oligomers were strongly increased relative to wild-type *App^NL-F/NL-F^* mouse brain levels, in the absence of an effect on APP precursor or monomeric Aβ levels. No *Sst* gene- associated effect on tau protein levels or phosphorylation were observed.

This study draws into question widely held concepts regarding the dominant mechanism by which Sst influences Aβ deposition in AD. For a more nuanced interpretation, it is critical to consider prior pertinent data. In 2005, a landmark study based on the same *Sst* knockout line employed in this work reported that neprilysin protein levels and activity were lower in the hippocampi, but not the cortices or cerebella, of *Sst* null mice with a corresponding increase in Aβ levels restricted to the hippocampus (26). The authors proposed a model whereby the effect of the *Sst* gene knockout on Aβ levels is mediated by the previously established ability of neprilysin to digest Aβ; in other words, it was concluded that Sst acts indirectly on Aβ by inducing the expression of neprilysin. No data on Aβ aggregation were generated at the time because the mice employed only produce endogenous levels of mouse Aβ, and even a full knockout of the Aβ-degrading enzyme neprilysin does not generate Aβ amyloidosis in this model (45). Nonetheless, the authors proposed that this phenomenon might also explain how diminishing SST levels, which have long been known to drop faster in late-onset AD cases relative to age-matched healthy elderly individuals (16, 17, 46, 47), may contribute to the disease. More recently, the main findings of this earlier study were corroborated, as repeated injections of Sst recombinantly fused to a blood brain barrier transporter (SST-scFv8D3) increased hippocampal neprilysin expression and reduced membrane-bound hippocampal Aβ_42_ in transgenic mice engineered to overexpress human APP harboring the Swedish mutation (48). Although the use of an AD mouse model could have lent itself to observing whether SST modulated Aβ aggregation, this was not investigated.

The present work revealed whole brain neprilysin levels to be unaffected by *Sst* ablation at the mRNA or protein levels. It could be said therefore that a caveat in the interpretation of these divergent results is the difference in brain structures analyzed, i.e., hippocampus versus whole brain, including neprilysin-rich structures (basal ganglia, pituitary), which could dilute hippocampus-specific effects. Consistent with this interpretation, the difference in plaque numbers that we observed at 12 months were more pronounced in the hippocampus than cortical areas. More specifically, only in the hippocampi of 12-month-old mice did any plaque size range (plaques with areas 100.1-200 square micrometers) satisfy the t-test 0.05 confidence threshold. As we did not survey RNA expression in the hippocampus specifically, we cannot reject the hypothesis that this Sst-dependent local inhibition of plaque formation is due to a functional interaction between Sst and neprilysin. That said, according to recent Human Protein Atlas data (https://www.proteinatlas.org/ENSG00000196549-MME/brain) the hippocampus expresses a small fraction of total brain neprilysin mRNA and protein, such that if hippocampal neprilysin levels were to increase several-fold while remaining constant elsewhere, total expression would still be subtly affected (49). Therefore, one conclusion of our study is that Sst is likely to play a lesser overall role for the expression of neprilysin throughout the brain than was previously appreciated.

We are not the first to suggest that the relationship between Sst and neprilysin expression may be limited to specific brain structures. For instance, the initial report establishing this relationship noted that in the cerebellum neprilysin is insensitive to *Sst* gene ablation (26). More recently, a reduction in the levels of human SST and CORT in the temporal lobe of AD postmortem brains could not be correlated with neprilysin levels (17). Finally, the aforementioned administration of the Sst-fusion construct only changed neprilysin levels in the hippocampus (48).

How else, if not on the basis of SST controlling whole brain neprilysin expression, can we account for our observation that SST ablation increases the proportion of small plaques? We previously showed that Sst binds to oligomeric Aβ and modulates Aβ aggregation *in vitro*, preventing the formation of amyloid fibrils that can incorporate thioflavin T (31). A closer look at fractions generated by co-incubation of Aβ and Sst showed that Sst delayed Aβ aggregation by stabilizing smaller oligomeric Aβ structures. More recently, a separate group reported that SST can bind certain Aβ oligomers (32). Consistent with these earlier data, the *in vivo* immunohistochemical observations reported here support the notion that Sst inhibits early events in amyloid plaque formation such that plaques form more readily in *Sst* knockouts, then grow at a rate unaffected by *Sst* genotype (**Fig 6**).

**Figure 6:**
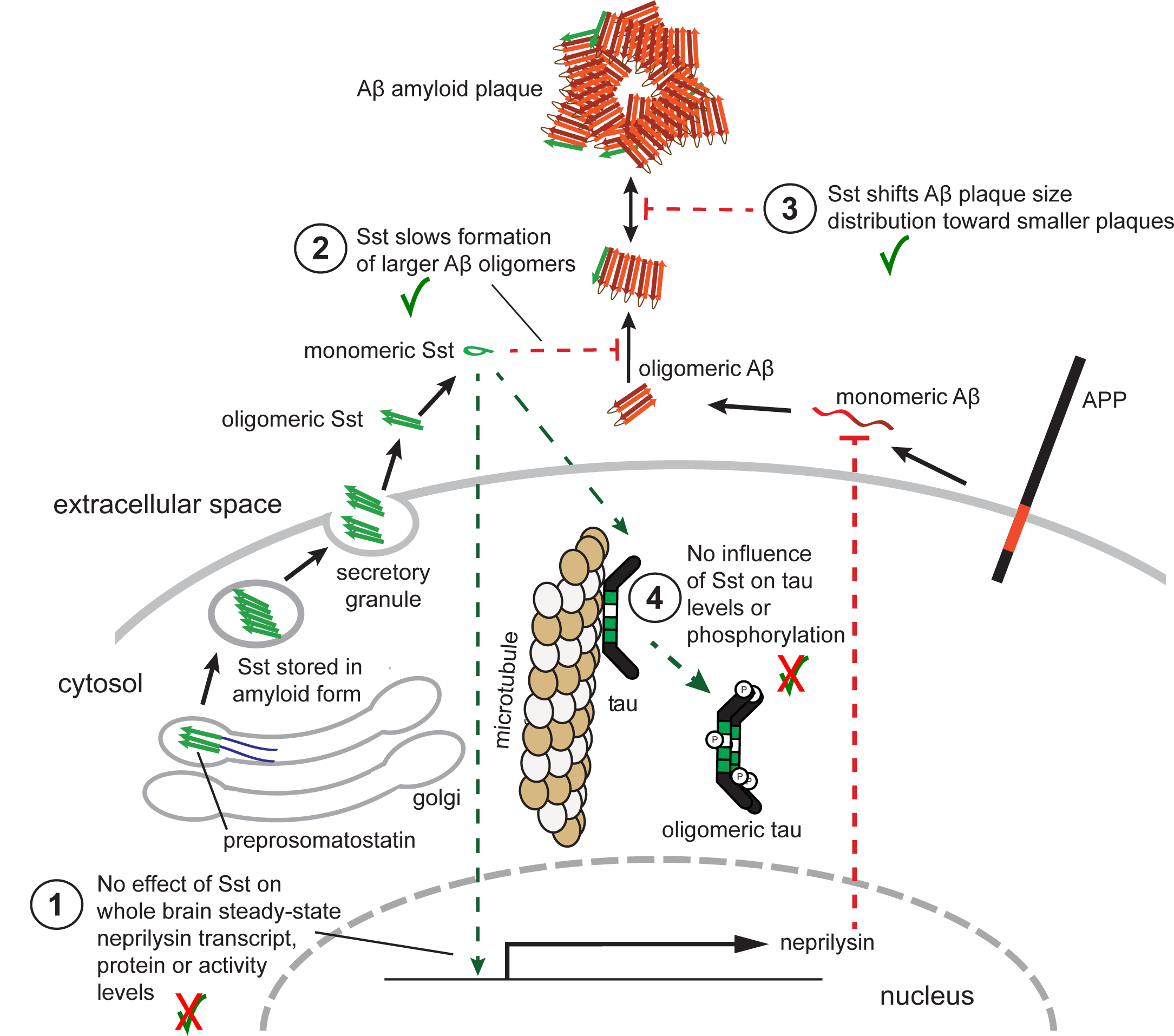
Cartoon summarizing key findings of this study. In 15-month-old *App^NL-F/NL-F^*mice, the presence or absence of Sst has no effect on whole-brain steady-state neprilysin transcript, protein or activity levels (1). Instead, Sst slows the formation of Aβ oligomers that can be visualized as 55-150 kDa SDS-resistant 82E1-reactive bands (2) and shifts Aβ amyloid formation and deposition (3). No effect of Sst was observed on steady-state tau protein levels or the tau phosphorylation at key AD phospho-acceptor sites. This cartoon was adapted and modified from a previously published figure (see Figure 2 in (12)).

Arguably, our most instructive data for mechanistic modelling come from the western blotting of brain extracts derived from 15-month-old *App^NL-F/NL-F^Sst^+/+^ and App^NL-F/NL-F^Sst^-/^*^-^ mice. These western blot data established that the presence of Sst reduced the levels of 55-150 kDa bands that were reactive to an Aβ-specific antibody. That these signals originated from Aβ oligomers, as opposed to another APP isoform or cleavage product, can be deduced from the specificity of the antibody used, which is known to selectively target the N-terminal cleavage site of Aβ. Less obvious is whether the Sst-dependent slowing of Aβ aggregation would be therapeutic or detrimental for individuals with AD. Although it may be more intuitive to interpret the role of Sst as beneficial, whether this is true will depend on where exactly in the Aβ cascade Sst interferes, i.e., whether the presence of Sst increases or decreases the levels of the most toxic Aβ aggregation intermediates.

Could the *Sst* paralog *Cort* have compensated for the loss of *Sst* in this Aβ amyloidosis paradigm? This possibility is not far-fetched: Our prior *in vitro* data established that both Sst and Cort can interfere with Aβ aggregation, with Cort being the more potent inhibitor in the respective thioflavin T fluorescence assay (31). Moreover, a compensatory increase in hypothalamic *Cort* mRNA was documented in male *Sst* knockout mice (42). Consistent with this previous report, we also found *Cort* transcript levels to increase in *Sst* null mice by around 35%. In contrast to the prior work, however, we observed no difference in this regard between male and female *App^NL-F/NL-F^Sst^-/-^*mice. This apparent distinction may merely reflect our inclusion of the cerebrum, cerebellum, and caudoputamen, areas of the brain that express *Cort* and perhaps obscured sex-specific differences that may still exist in the hypothalamus.

When comparing female and male *APP^NL-F/NL-F^* mice of each of the three *Sst* genotypes, we observed no sex-specific differences in cortical and hippocampal plaque density and plaque size distributions. In the absence of a sex-specific phenotype, which could have clarified the possible influence of a compensatory biology, our data cannot resolve the extent to which *Cort*- based compensation may have precluded the formation of more striking differences in AD-like Aβ amyloidosis. A *Cort* null mouse (50) and an *Sst/Cort* double knockout mouse (51) have been reported, raising the possibility that a triple transgenic line could be produced.

Another factor that could have masked the physiological influence of Sst on Aβ amyloidosis and AD molecular signatures in this study is the artificially high level of Aβ_1-42_ that the *App^NL-F/NL-F^*knock-in mice are known to produce. This caveat represents a conceptual catch- 22 because the very ability to measure the influence of Sst on Aβ amyloidosis depends on a paradigm that can produce sufficient Aβ levels to study its aggregation. Thus, any anticipation that the ablation of endogenous Sst would by itself profoundly affect amyloid plaque formation in *APP^NL-F/NL-F^*mice needs to be tempered by the recognition that Aβ deposition in this model depends on Aβ_1-42_ production that far exceeds physiological Aβ production in human AD brains. When considered in this context, it is remarkable that *Sst* gene ablation alone was sufficient to increase the levels of SDS-stable Aβ assemblies to the extent seen.

## CONCLUSIONS

The results of this study document that Sst does not alter steady-state neprilysin transcript, protein or activity levels in whole brain extracts yet represent robust *in vivo* evidence of an inverse relationship between Sst levels and Aβ deposition. As such, the results are easier reconciled with a model of Sst affecting Aβ aggregation directly, consistent with recent biochemical and molecular dynamics simulation studies. This scenario also fits with results from new work documenting a relatively close spatial correlation of Sst and Aβ release sites in various brain regions (52). Future work will need to establish if the combined ablation of the *Sst* and *Cort* genes will further enhance the impact on Aβ amyloid deposition. Once we understand if the Sst-mediated slowing of Aβ aggregation reduces or exacerbates toxicity *in vivo*, refined strategies should come to the fore that can harness the still untapped therapeutic potential of this cyclic neuropeptide for the treatment of AD.

## ACKNOWLEDGEMENTS

The authors thank Dr. Paul McKeever, University of Toronto, for highly informative discussions on immunohistochemistry and neuroanatomy, Zhilan Wang, University of Toronto, for tissue embedding, sectioning and slide mounting, and Erica Stuart, University of Toronto, for help with RT-qPCRs.

## SUPPORTING INFORMATION

S1 Figure. Indel sequence of *Sst^-/-^* allele determined by Sanger sequencing.

In addition to depicting the indel, the image shows the position of forward and reverse primers used in this study for genotyping.

S2 Figure. Hippocampal Aβ amyloid plaque counts in 12- or 15-month-old *App^NL-F/NL-F^* mice that express wild-type Sst levels or were Sst-deficient.

Hippocampal Aβ amyloid plaque densities increased between 12- and 15-month-old *App^NL-F/NL-F^* mice. Differences in hippocampal Aβ amyloid plaque densities were observed when comparing *App^NL-F/NL-F^*mice that expressed wild-type Sst versus Sst-deficient mice. More specifically, a trend toward higher Aβ plaque densities of small sizes (0.25-200 µm) was observed in 12- month-old *Sst* gene-deficient mice, echoing the increase in Aβ amyloid plaque densities observed in the cortex of *Sst* ablated *App^NL-F/NL-F^* mice (Fig 3E).

## Notes

### Competing Interest Statement

The authors have declared no competing interest.

## REFERENCES

1. Vale W, Brazeau P, Grant G, Nussey A, Burgus R, Rivier J, et al. [Preliminary observations on the mechanism of action of somatostatin, a hypothalamic factor inhibiting the secretion of growth hormone]. Comptes rendus hebdomadaires des seances de l’Academie des sciences Serie D: Sciences naturelles. 1972;275(25):2913–6.

2. Brazeau P, Vale W, Burgus R, Ling N, Butcher M, Rivier J, et al. Hypothalamic polypeptide that inhibits the secretion of immunoreactive pituitary growth hormone. Science. 1973;179(4068):77-9.

3. de Lecea L, Criado JR, Prospero-Garcia O, Gautvik KM, Schweitzer P, Danielson PE, et al. A cortical neuropeptide with neuronal depressant and sleep-modulating properties. Nature. 1996;381(6579):242-5.

4. Martel G, Dutar P, Epelbaum J, Viollet C. Somatostatinergic systems: an update on brain functions in normal and pathological aging. Frontiers in Endocrinology. 2012;3(154):1–15.

5. Patel YC, Greenwood MT, Warszynska A, Panetta R, Srikant CB. All five cloned human somatostatin receptors (hSSTR1-5) are functionally coupled to adenylyl cyclase. Biochem Biophys Res Commun. 1994;198(2):605–12.

6. Theodoropoulou M, Stalla GK. Somatostatin receptors: from signaling to clinical practice. Frontiers in neuroendocrinology. 2013;34(3):228–52.

7. Siehler S, Nunn C, Hannon J, Feuerbach D, Hoyer D. Pharmacological profile of somatostatin and cortistatin receptors. Mol Cell Endocrinol. 2008;286(1-2):26–34.

8. Deghenghi R, Papotti M, Ghigo E, Muccioli G. Cortistatin, but not somatostatin, binds to growth hormone secretagogue (GHS) receptors of human pituitary gland. Journal of endocrinological investigation. 2001;24(1):Rc1-3.

9. Dalm VA, Van Hagen PM, de Krijger RR, Kros JM, Van Koetsveld PM, Van Der Lely AJ, et al. Distribution pattern of somatostatin and cortistatin mRNA in human central and peripheral tissues. Clin Endocrinol (Oxf). 2004;60(5):625–9.

10. De Lecea L, del Rio, J.A., Criado, J.R., Alcantara, A., Morales, M., Danielson, P.E., et al. . Cortistatin Is Expressed in a Distinct Subset of Cortical Interneurons. J Neurosci. 1997;7(15):13.

11. Spier AD, de Lecea L. Cortistatin: a member of the somatostatin neuropeptide family with distinct physiological functions. Brain Res Brain Res Rev. 2000;33(2-3):228–41.

12. Solarski M, Wang H, Wille H, Schmitt-Ulms G. Somatostatin in Alzheimer’s disease: A new Role for an Old Player. Prion. 2018;12(1):1–8.

13. Epelbaum J, Guillou JL, Gastambide F, Hoyer D, Duron E, Viollet C. Somatostatin, Alzheimer’s disease and cognition: an old story coming of age? Prog Neurobiol. 2009;89(2):153–61.

14. Burgos-Ramos E, Hervas-Aguilar A, Aguado-Llera D, Puebla-Jimenez L, Hernandez-Pinto AM, Barrios V, et al. Somatostatin and Alzheimer’s disease. Mol Cell Endocrinol. 2008;286(1-2):104–11.

15. Rossor MN, Emson PC, Mountjoy CQ, Roth M, Iversen LL. Reduced amounts of immunoreactive somatostatin in the temporal cortex in senile dementia of Alzheimer type. Neurosci Lett. 1980;20(3):373–7.

16. Davies P, Katzman R, Terry RD. Reduced somatostatin-like immunoreactivity in cerebral cortex from cases of Alzheimer disease and Alzheimer senile dementa. Nature. 1980;288(5788):279–80.

17. Gahete MD, Rubio A, Duran-Prado M, Avila J, Luque RM, Castano JP. Expression of Somatostatin, cortistatin, and their receptors, as well as dopamine receptors, but not of neprilysin, are reduced in the temporal lobe of Alzheimer’s disease patients. J Alzheimers Dis. 2010;20(2):465–75.

18. Ramos B, Baglietto-Vargas D, del Rio JC, Moreno-Gonzalez I, Santa-Maria C, Jimenez S, et al. Early neuropathology of somatostatin/NPY GABAergic cells in the hippocampus of a PS1xAPP transgenic model of Alzheimer’s disease. Neurobiol Aging. 2006;27(11):1658–72.

19. Gaspar P, Duyckaerts C, Febvret A, Benoit R, Beck B, Berger B. Subpopulations of somatostatin 28-immunoreactive neurons display different vulnerability in senile dementia of the Alzheimer type. Brain Res. 1989;490(1):1–13.

20. Waller R, Mandeya M, Viney E, Simpson JE, Wharton SB. Histological characterization of interneurons in Alzheimer’s disease reveals a loss of somatostatin interneurons in the temporal cortex. Neuropathology. 2020;40(4):336–46.

21. Sos KE, Mayer MI, Takács VT, Major A, Bardóczi Z, Beres BM, et al. Amyloid β induces interneuron-specific changes in the hippocampus of APPNL-F mice. PLoS One. 2020;15(5):e0233700.

22. Roberts GW, Crow TJ, Polak JM. Location of neuronal tangles in somatostatin neurones in Alzheimer’s disease. Nature. 1985;314(6006):92–4.

23. Morrison JH, Rogers J, Scherr S, Benoit R, Bloom FE. Somatostatin immunoreactivity in neuritic plaques of Alzheimer’s patients. Nature. 1985;314(6006):90–2.

24. Vepsalainen S, Helisalmi S, Koivisto AM, Tapaninen T, Hiltunen M, Soininen H. Somatostatin genetic variants modify the risk for Alzheimer’s disease among Finnish patients. J Neurol. 2007;254(11):1504–8.

25. Xue S, Jia L, Jia J. Association between somatostatin gene polymorphisms and sporadic Alzheimer’s disease in Chinese population. Neurosci Lett. 2009;465(2):181–3.

26. Saito T, Iwata N, Tsubuki S, Takaki Y, Takano J, Huang SM, et al. Somatostatin regulates brain amyloid beta peptide Abeta42 through modulation of proteolytic degradation. Nat Med. 2005;11(4):434–9.

27. Iwata N, Tsubuki S, Takaki Y, Watanabe K, Sekiguchi M, Hosoki E, et al. Identification of the major Abeta1-42-degrading catabolic pathway in brain parenchyma: suppression leads to biochemical and pathological deposition. Nat Med. 2000;6(2):143–50.

28. Fleisher-Berkovich S, Filipovich-Rimon T, Ben-Shmuel S, Hülsmann C, Kummer MP, Heneka MT. Distinct modulation of microglial amyloid β phagocytosis and migration by neuropeptides. J Neuroinflammation. 2010;7:61.

29. Ciaccio C, Tundo GR, Grasso G, Spoto G, Marasco D, Ruvo M, et al. Somatostatin: a novel substrate and a modulator of insulin-degrading enzyme activity. J Mol Biol. 2009;385(5):1556–67.

30. Tundo G, Ciaccio C, Sbardella D, Boraso M, Viviani B, Coletta M, et al. Somatostatin modulates insulin-degrading-enzyme metabolism: implications for the regulation of microglia activity in AD. PLoS One. 2012;7(4):e34376.

31. Wang H, Muiznieks LD, Ghosh P, Williams D, Solarski M, Fang A, et al. Somatostatin binds to the human amyloid beta peptide and favors the formation of distinct oligomers. Elife. 2017;6:e28401.

32. Puig E, Tolchard J, Riera A, Carulla N. Somatostatin, an In Vivo Binder to Aβ Oligomers, Binds to βPFO(Aβ(1-42)) Tetramer. ACS Chem Neurosci. 2020;11(20):3358–65.

33. van Grondelle W, Iglesias CL, Coll E, Artzner F, Paternostre M, Lacombe F, et al. Spontaneous fibrillation of the native neuropeptide hormone Somatostatin-14. J Struct Biol. 2007;160(2):211–23.

34. Maji SK, Perrin MH, Sawaya MR, Jessberger S, Vadodaria K, Rissman RA, et al. Functional amyloids as natural storage of peptide hormones in pituitary secretory granules. Science. 2009;325(5938):328–32.

35. Anoop A, Ranganathan S, Das Dhaked B, Jha NN, Pratihar S, Ghosh S, et al. Elucidating the role of disulfide bond on amyloid formation and fibril reversibility of somatostatin- 14: relevance to its storage and secretion. J Biol Chem. 2014;289(24):16884–903.

36. Saito T, Matsuba Y, Mihira N, Takano J, Nilsson P, Itohara S, et al. Single App knock-in mouse models of Alzheimer’s disease. Nat Neurosci. 2014;17(5):661–3.

37. Zeyda T, Diehl N, Paylor R, Brennan MB, Hochgeschwender U. Impairment in motor learning of somatostatin null mutant mice. Brain Res. 2001;906(1-2):107–14.

38. Livak KJ, Schmittgen TD. Analysis of relative gene expression data using real-time quantitative PCR and the 2(-Delta Delta C(T)) Method. Methods. 2001;25(4):402–8.

39. Zeyda T, Hochgeschwender U. Null mutant mouse models of somatostatin and cortistatin, and their receptors. Mol Cell Endocrinol. 2008;286(1-2):18–25.

40. Wang H, Williams D, Griffin J, Saito T, Saido TC, Fraser PE, et al. Time-course global proteome analyses reveal an inverse correlation between Abeta burden and immunoglobulin M levels in the APPNL-F mouse model of Alzheimer disease. PLoS One. 2017;12(8):e0182844.

41. Aladeokin AC, Akiyama T, Kimura A, Kimura Y, Takahashi-Jitsuki A, Nakamura H, et al. Network-guided analysis of hippocampal proteome identifies novel proteins that colocalize with Aβ in a mice model of early-stage Alzheimer’s disease. Neurobiol Dis. 2019;132:104603.

42. Luque RM, Kineman RD. Gender-dependent role of endogenous somatostatin in regulating growth hormone-axis function in mice. Endocrinology. 2007;148(12):5998–6006.

43. Wu M, Dorosh L, Schmitt-Ulms G, Wille H, Stepanova M. Aggregation of Aβ40/42 chains in the presence of cyclic neuropeptides investigated by molecular dynamics simulations. PLoS Comput Biol. 2021;17(3):e1008771.

44. Horikoshi Y, Mori T, Maeda M, Kinoshita N, Sato K, Yamaguchi H. Abeta N-terminal-end specific antibody reduced beta-amyloid in Alzheimer-model mice. Biochem Biophys Res Commun. 2004;325(2):384–7.

45. Iwata N, Tsubuki S, Takaki Y, Shirotani K, Lu B, Gerard NP, et al. Metabolic regulation of brain Abeta by neprilysin. Science. 2001;292(5521):1550–2.

46. Saiz-Sanchez D, Ubeda-Banon I, de la Rosa-Prieto C, Argandona-Palacios L, Garcia- Munozguren S, Insausti R, et al. Somatostatin, tau, and beta-amyloid within the anterior olfactory nucleus in Alzheimer disease. Exp Neurol. 2010;223(2):347–50.

47. Lu T, Pan, Y., Kao, S.Y., Li, C., Kohane, I., Chan, J., and Yanker, B.A. Gene regulation and DNA damage in the ageing human brain. Nature. 2004;429(6994):9.

48. Rofo F, Ugur Yilmaz C, Metzendorf N, Gustavsson T, Beretta C, Erlandsson A, et al. Enhanced neprilysin-mediated degradation of hippocampal Aβ42 with a somatostatin peptide that enters the brain. Theranostics. 2021;11(2):789–804.

49. Uhlén M, Fagerberg L, Hallström BM, Lindskog C, Oksvold P, Mardinoglu A, et al. Proteomics. Tissue-based map of the human proteome. Science. 2015;347(6220):1260419.

50. Luque RM, Cordoba-Chacon J, Pozo-Salas AI, Porteiro B, de Lecea L, Nogueiras R, et al. Obesity- and gender-dependent role of endogenous somatostatin and cortistatin in the regulation of endocrine and metabolic homeostasis in mice. Sci Rep. 2016;6(1):37992.

51. Pedraza-Arevalo S, Cordoba-Chacon J, Pozo-Salas AI, F LL, de Lecea L, Gahete MD, et al. Not So Giants: Mice Lacking Both Somatostatin and Cortistatin Have High GH Levels but Show No Changes in Growth Rate or IGF-1 Levels. Endocrinology. 2015;156(6):1958–64.

52. Watamura N, Kakiya N, Nilsson P, Tsubuki S, Kamano N, Takahashi M, et al. Somatostatin-evoked Aβ catabolism in the brain: Mechanistic involvement of α- endosulfine-K(ATP) channel pathway. Mol Psychiatry. 2021;Epub:DOI: 10.1038/s41380-021-01368-8.

