## Supplementary figures and images for "Somatostatin slows Aβ plaque deposition in aged *APP^NL-F/NL-F^* mice by blocking Aβ aggregation in a neprilysin-independent manner"

### Supporting Information

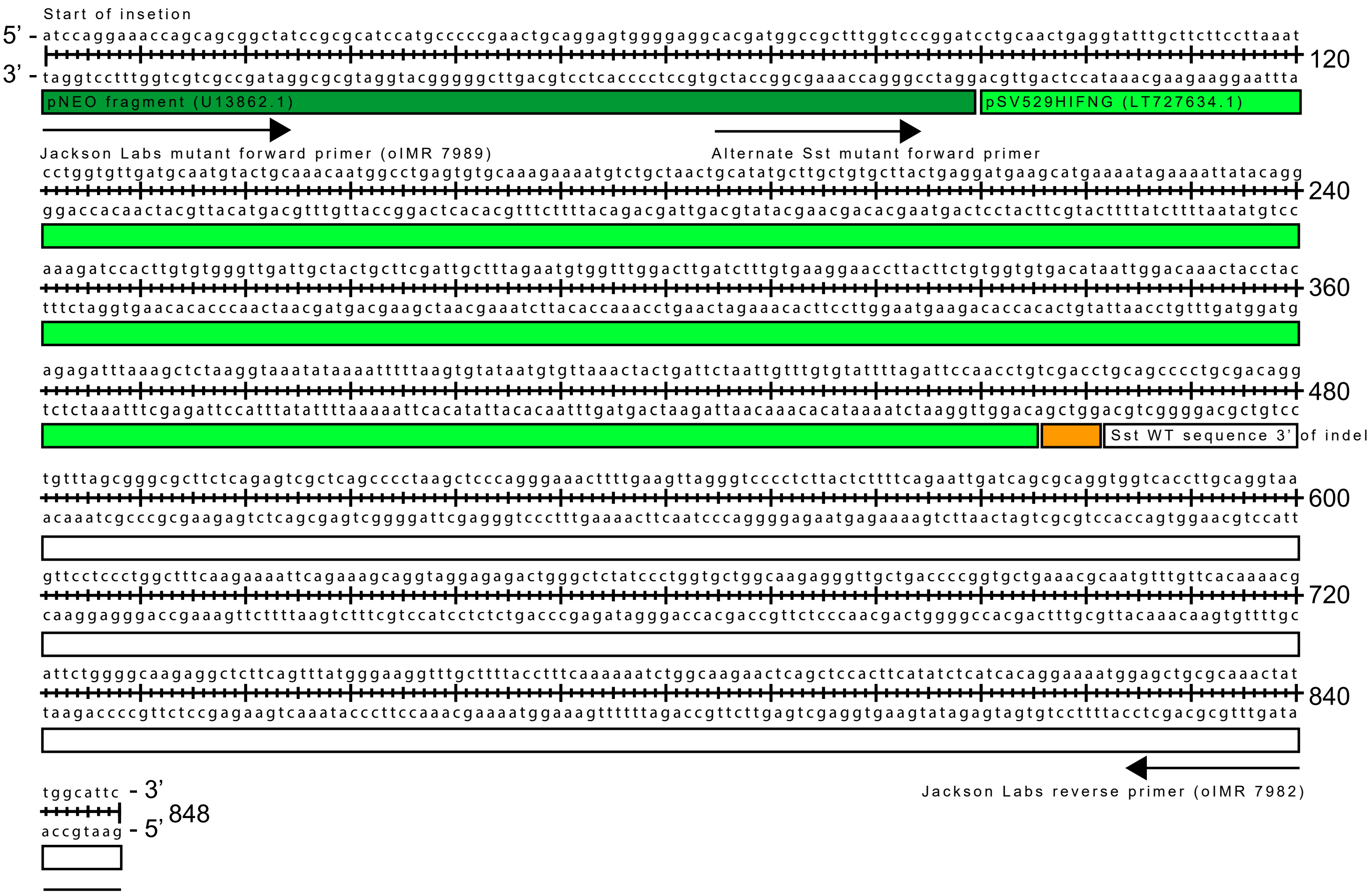

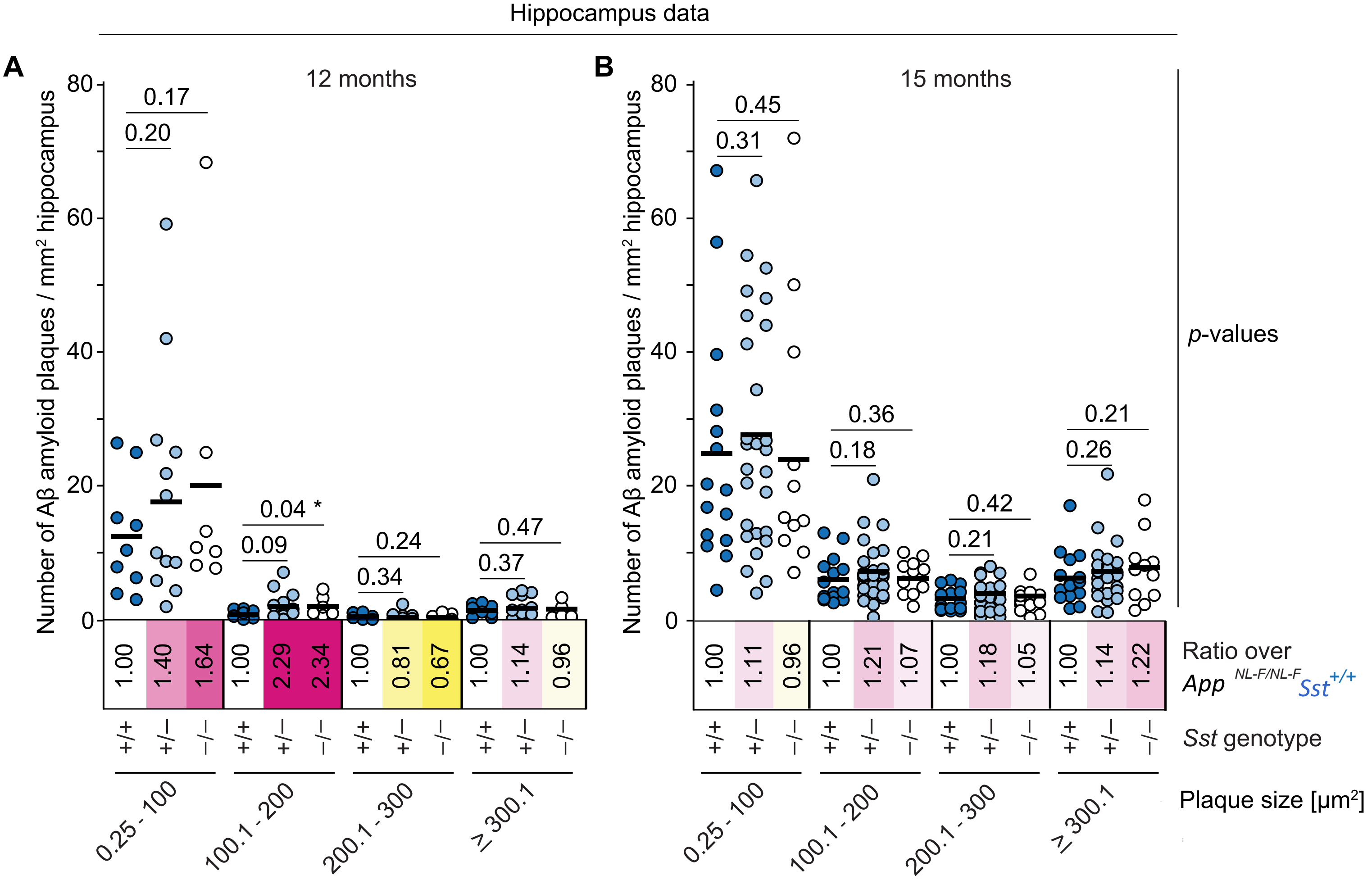
